# Multivalent Adhesive Probe Atomic Force Microscopy (MAPA) for accessing dispersive adhesion of cells and biosurfaces

**DOI:** 10.64898/2026.08.31.748212

**Authors:** Maria Gaczynska, Pawel A. Osmulski

## Abstract

Adhesion of cells is the key factor determining functioning of multicellular organisms. Viscoelastic properties of cells can be studied by multiple methods. However, attractiveness of cells or extracellular matrix without the elastic component (dispersive adhesion) is not accessible. We present an extension of force spectrometry technology: the Multivalent Adhesive Probe Atomic Force Microscopy (MAPA) that delivers dispersive adhesion maps of live cells and biosurfaces, and identifies differences unresolved by viscoelastic probing.

---

Interaction of cells with their environment is achieved by specific and nonspecific forces including cohesive and adhesive components amounting for cell “attractiveness”: the major biophysical property determining the cell fate and crucial for organ development as well as for cancer invasiveness. However, physical parameters are rarely used for cell characterization despite their exceptional sensitivity to physiological status, often early-pointing to the changes undetectable by biochemical or genetic methods^1,2,3^. Cell elasticity is the most often studied physical property^4,5^. Another parameter amounting for the “biophysical phenotype” of cancer cells and crucial for tumor invasion is surface adhesion^6^. Formally, adhesion is defined as the free energy change to separate unit areas of two media from contact to infinity. The cell adhesiveness is a result of an interplay of nonspecific physical and chemical properties of surface biomolecules^7–9^ and specific interactions of surface receptors, the latter studied extensively in the context of cancer^10–13^, the former often overlooked. Methods to study adhesiveness are either non-quantitative or access viscoelastic adhesion, or target specific surface receptors^14^ (see Methods). In force spectrometry (FS) – AFM used to access viscoelastic adhesion a micrometer-sized tip on a cantilever (the probe) raster scans the cell’s surface with indentations at each cell-tip contact. Viscoelastic adhesion offers important insight into cell’s propensity to establish stable binding through adhesion molecules and surface receptors. However, the first contact between cells or cells and ECM may be governed by van der Waals interactions based dispersive adhesion. Dispersive adhesion is likely a critical first step for enabling (or preventing) the subsequent stable binding. However, dispersive adhesion of cells is not accessible by known biophysical methods.

Here, we present a novel extension of the atomic force microscopy (AFM) technology to compare the dispersive adhesion of living cells or other surfaces, without the elastic component of the object indentation by the FS-AFM. The Multivalent Adhesive Probe Atomic Force Microscopy (MAPA) relies on a naturally multivalent combination of London, Debye and Keesom van der Walls (VDW) forces^15^. Additional “multivalency” is offered by probe modifications with chemical entities of choice. We took inspiration from a non-contact oscillation mode AFM, where the probe oscillates with or near a fundamental resonant frequency above the surface and is kept within the VDW distance when creating the topographical image of the (usually) nano-sized surface. For biologically relevant probing in liquid, the fundamental resonant frequency is low (9-10 kHz), and the tuning spectrum recorded without any contact with the sample (non-engaged probe) is rich in harmonic frequencies and “parasitic” frequencies, and some thermal noise. All these unique eigenfrequencies other than single fundamental resonant frequency are neglected under the standard use of non-contact AFM. Instead of exciting the probe with a single eigenfrequency, we record a broad-range (0 kHz to up to 800 kHz) tuning spectrum A=f(F) (amplitude A as a function of frequency F) of cantilever vibrations. Moreover, we record the spectrum not only when the cantilever is hovering high (micrometers) above the probed surface (the “substrate”), as practiced for finding the single resonant frequency, but also when the AFM probe is at the VDW distance from the substrate, engaged in VDW interactions (**Fig. 1A**). The probed surface can be a cell or other biological, organic or inorganic entity. We base our method on the observation that the spectra of a non-engaged probe differ from the engaged probe spectra **(Fig. 1B)**. We tested the method first with model coated substrates representing a bare chemically inert mineral (muscovite mica), a coat of amine-rich polymer polyethyleneimine (PEI), a protein (bovine serum albumin; BSA) and a lipid dipalmitoyl D-α-phosphatidylcholine (DPPC) (**Fig. 1C**). The cantilever and tip were excited bare (chemically inert silicon nitrate; SiN) or coated with BSA or PEI (**Fig. 1C-H**). Understandably, non-engaged spectra reflect the type of a probe coating (**Fig. 1D**), whereas engaged probes tend to agglomerate according to substrate chemical properties (**Fig. 1E**). The engaged spectra change depending specifically on the physicochemical characteristics of the probe and the substrate (**Fig. 1F-H**). In essence, by analysis of engaged-probe spectra we gain insight into organization of dipole moments on the probed surfaces.

**Fig. 1.**
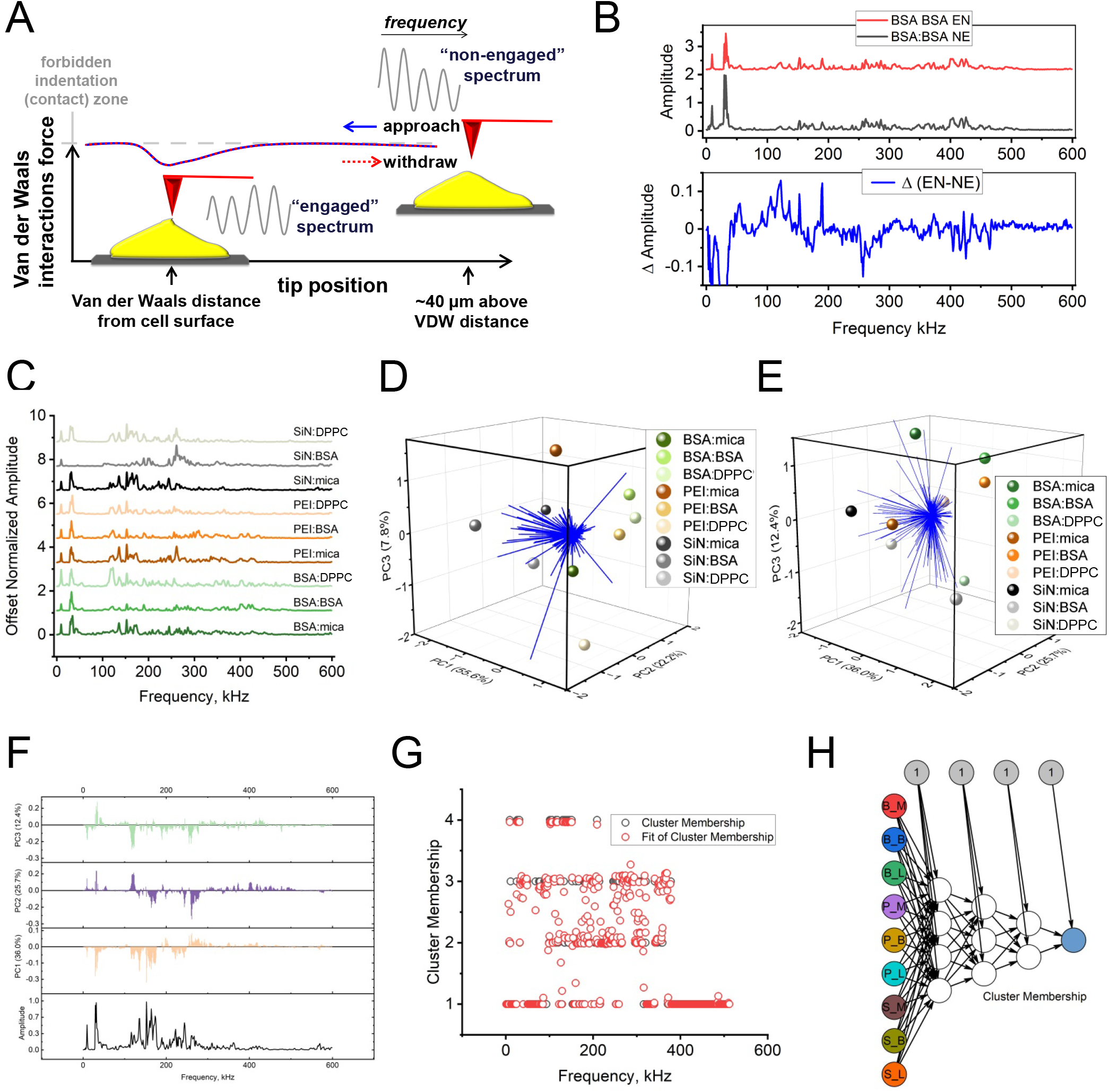
Probing of model surfaces with Multivalent Adhesive Probe AFM (MAPA). **A**.Schematics of MAPA. The probes (red) were tuned to near-resonant fundamental frequency in liquid (∼ 9 kHz) and engaged with the cell or model surface with negligibly small scanning area. After examining the force curve and the Z-position of probe to assure that that engagement is not “false” and that only VDW interactions are present (no surface indentation in “forbidden “zone), the sweep of 0 to up to 800 kHz frequencies was applied to the probe and the “engaged” frequency spectra were collected. Then, after return to scanning and checking the proper VDW probe engagement, the probe was lifted about 40 µm and the “non-engaged” reference spectra were collected. The spectra were processed and analyzed as presented below and described in Methods. **B**.Examples of spectra collected consecutively with BSA-coated probe interrogating BSA-coated mica: “engaged” (top/red; EN), “non-engaged” (middle/black; NE) and a difference between them (bottom; Δ). **C – H**. Spectral analysis reveals separation of “signatures” characterizing model surfaces. **C**. Stacked engaged spectra. Here and in subsequent legends probe coating is listed first, substrate coating is listed second. 3D Biplots of spectral Principal Component Analysis (PCA) performed on amplitude spectra produce separate populations of scores and loads obtained with: **D**. non-engaged and **E**. engaged probes. Spheres represent scores and blue vectors – loadings (frequencies). **F**. Loading plots of spectral PCA showing frequencies that distinguish between probe status and different substrate coatings. Spectra of uncoated tip and mica were used as a reference (example of the spectrum at the bottom). **G**. Results of Neural Network classification of engaged spectra using the resilient backpropagation with back tracking algorithm. Classification power was R=0.86. **H**. Graph Neural network of surface recognition. See Methods for details.

After assessing interaction between model organic and inorganic surfaces of the probe and substrate, we turned into live cultured model prostate cancer cells, 22Rv1. Instead of relying on cell’s indentation, we probed them with MAPA, with SiN (uncoated) probes or probes coated with BSA. **Fig. 2A,B** presents results of spectral analysis of the engaged spectra collected when a cell was the substrate, or with mica or PEI coated substrates. The heat map in **Fig. 2A** shows distinct clustering of frequencies that may constitute selection of analytical indicators distinguishing between substrates (cells or model surfaces) or coatings. Interrogation of model substrates resulted in uniform sets of spectra without classes distinguished for mica or PEI categories. To the contrary, properties of the 22Rv1 cells allowed distinguishing two classes each within the categories of cells interrogated with uncoated or BSA-coated probes (**Fig. 2A**). Further machine learning supported classification of spectra generated with cells revealed four clusters (**Fig. 2B**). We may speculate that MAPA senses properties of cells not obvious by assessment of the cells’ morphology, always performed before AFM probing.

**Fig. 2.**
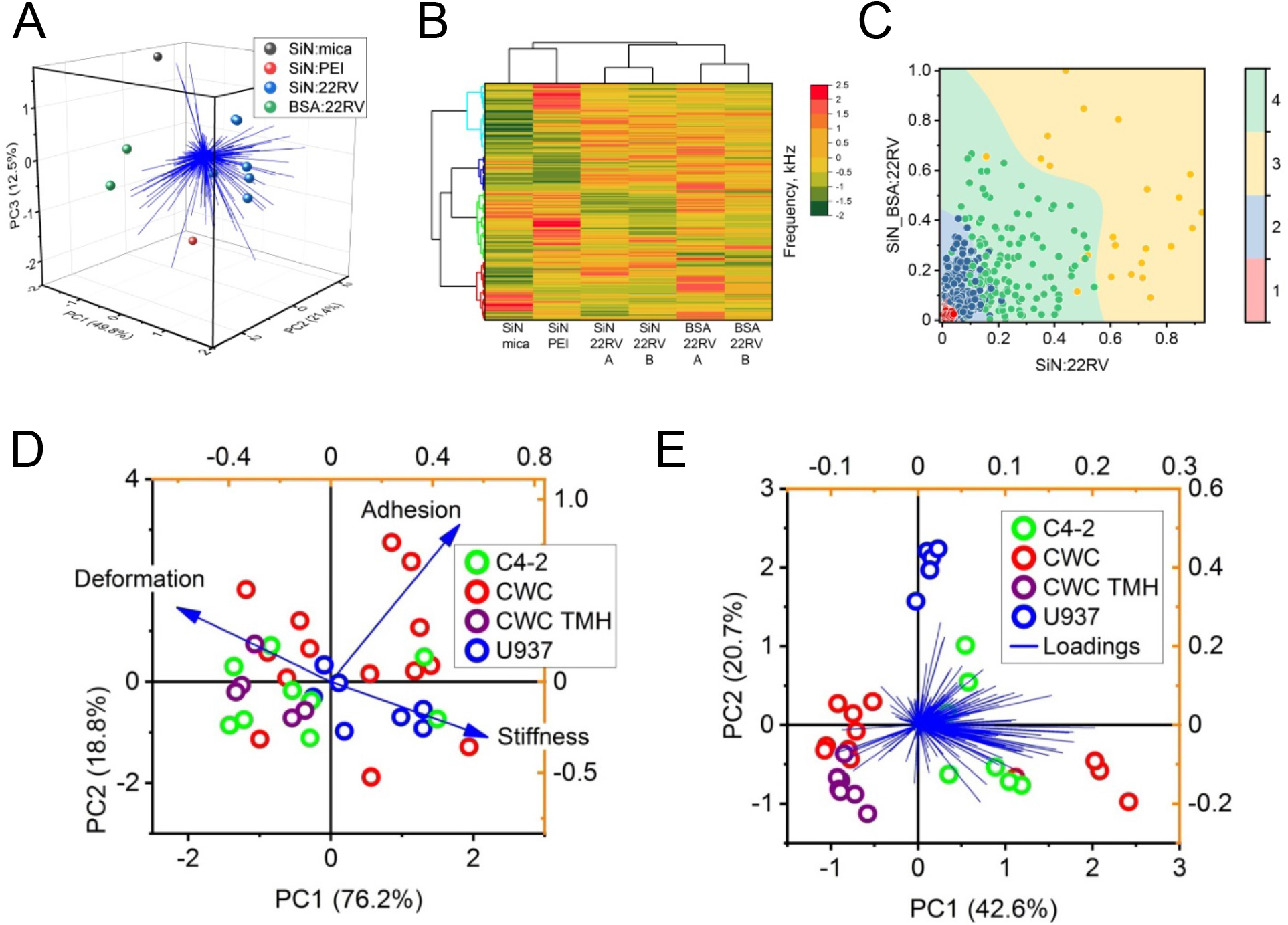
Distinct properties of live human cells are accessible to MAPA probing. **A-C**: 22Rv1 cultured model prostate cancer cells. **A**. 3D spectral PCA separates properties of engaged spectra depending on the applied probe and tested substrate. **B**. Heat map shows distinct clustering of frequencies that may constitute selection of analytical indicators. Each column represents means of 2-3 spectra. **C**. Support vector machine (SVM) classification with the RBF kernel of spectra of 22RV1 cells obtained with uncoated and BSA doped probes. Clusters were identified with accuracy rate 0.804. One dot = one frequency. **D, E**. Cultured model prostate cancer C4-2 cells (eGFP), U937 derived macrophages (mCherry) and CWC: a stable tumor-macrophage hybrid (TMH) cell line formed from C4-2 and U937 macrophages, as described. ^16^ CWC TMHs are fusions newly generated in the co-culture of CWC and C4-2 and identified by strong eGFP and mCherry signals (CWC: weak eGFP, strong mCherry). **D**. PCA of mechanical parameters (loadings): stiffness, deformation and viscoelastic adhesion collected with indentation-based PF-QNM and **E**. Spectral PCA performed on engaged amplitude spectra collected with MAPA (frequencies as loadings). In both D and E one dot = one cell. Cells were captured on PEI-glass.

Encouraged by the diversification of seemingly homogenous population of cultured cells, we tested a set of model prostate cancer cells C4-2, model macrophages derived from U937 human monocytes, and CWC, a stable line of tumor-macrophage hybrids (TMHs) created *in vitro* by co-culture of C4-2 and U937 derived macrophages^16^. CWC cells co-cultured with C4-2 cancer cells display strong phagocytic properties; CWCs that based on microscopic evaluation just phagocytosed a cancer cell were labeled as CWC-TMH. In parallel with MAPA evaluation we collected mechanical phenotypes for the same cell types using Peak Force Quantitative Nanomechanics, an advanced type of FS-AFM that relies on cells’ indentation to collect data on viscoelastic adhesion, stiffness, and deformation. Apparently, PCA analysis of PF-QNM based data was not able to separate cancer cells from macrophages or hybrids (**Fig. 2C**). However, PCA based on MAPA-derived spectra did produce clearly separated classes of cells. Interestingly, CWC were separated into two classes, one of them clustering with CWC-TMH (**Fig. 2D**). We may speculate that these two groups may indicate separation of invasive and phagocytic propensities of CWCs, the latter common with actively phagocytosing CWC-TMHs, the former unique.

**In summary**, the method offers a unique insight into the propensity of biological objects to engage in interactions and into the chemical make-up of biological surfaces. We plan to expand MAPA into semi-quantitative or quantitative assessment of dispersive adhesion, especially for the purpose of comparison of surfaces. A detailed analysis of chemical composition will also be developed. The method will help to answer a broad array of biological questions on cells’ function and integrity.

## Methods

MAPA offers the first available method to assess dispersive adhesion of biological objects. Despite the significance of cell adhesiveness, options to study the phenomenon are limited. Available methods are either nonspecific and non-quantitative (scratch test for cell motility, shake-off test, enumeration of cell clusters, centrifugal assays, coating with fluorescent particles) or deliver data on viscoelastic properties (micropipette manipulations, microfluidic channels, atomic force microscopy /AFM-based force spectrometry)^14,17^. Peak Force Quantitative Nanomechanical (PF-QNM) AFM delivers highly sensitive quantitative viscoelastic-adhesiveness data^11,12,18,19^. Most recent AFM-based nanoscale rheology accesses viscoelasticity as well^20^. In turn, chemical force AFM produces data only on the specific contribution of a selected single, component of the surface^21^. To the best of our knowledge, the idea of utilizing broad-spectrum frequencies for probing the dispersive adhesion has no precedent. Multi-frequency methods such as bimodal and “nanomechanical spectroscopy” AFMs are used to analyze elastic properties and composition of materials, including cell membranes^22–24^. Changes in resonant harmonic frequency of cantilever vibrations can be used to sense stiffness of materials^25^. In turn, ringing mode AFM offers compositional and adhesion mapping of cells’ or biomaterials’ surface, however it relies on cell indentation for viscoelastic properties and operates in air with fixed cells^26,27^.

Spectral analysis of oscillation AFM derived data that is in the center of MAPA is amenable to machine learning and general AI – supported upgrades. Provided below is the description of methods used so far to demonstrate the power of dispersive adhesion in differentiating between surfaces, model or cellular.

### Preparation of samples and probes

Model surfaces (AFM substrates) were prepared as follows: muscovite mica (model of chemically inert inorganic surface; Ted Pella Inc.; Redding, CA) was freshly cleaved and covered with PBS within 10 seconds. Branched polyethyleneimine (PEI; Sigma-Aldrich, Burlington, MA), a synthetic polymer with repeating amine groups and two-carbon aliphatic spacers, delivered an amine-rich coat prepared by covering freshly cleaved mica with 0.1% PEI in PBS, incubating for 10 minutes at room temperature (RT) then washing three times and then covering with PBS. Such prepared PEI-covered mica was also used to produce bovine serum albumin (BSA; Sigma-Aldrich, Burlington, MA) - coated substrates by covering the substrates with 1 mg/ml BSA in PBS for 10 min at RT, then washing three times and then covering with PBS. These substrates were labeled as “BSA coated”. Whenever practical, the same substrate was examined bare, PEI-coated and PEI-then-BSA coated in a sequence. The same procedure was used to coat probes with PEI and BSA, and to coat coverslip glass with PEI. Dipalmitoyl D-α- phosphatidylcholine (DPPC; Sigma-Aldrich, Burlington, MA) was dissolved in chloroform (5 mM), deposited on mica, evaporated under a stream of nitrogen and washed with PBS. We deliberately did not heat the lipid in order to obtain a tightly packed and rigid lipid deposit for point-probing, rather than water-penetrable fluid phase^28^. Cultured cells were attached to PEI-coated mica or glass by incubating cells’ suspension in PBS for 30 min in cell culture incubator, then washing three times with PBS. The following cell lines from ATCC (Manassas, VA) were cultured according to ATCC recommendations and used for MAPA probing: prostate cancer 22Rv1 (CRL-2505) and C4-2 (CRL-3314), and macrophages polarized from monocytoidal U937 (CRL-1593.2)^12^. CWC line: stable hybrids of C4-2 and U937-derived macrophages, were created and cultured as described in Chou et al.^16^. C4-2 and U937 were stably fluorescently labeled by introducing eGFP (green fluorescent protein) or mCherry (red; U937), to enable their identification in co-culture. Hybrid CWC cells displayed both eGFP and mCherry labels^16^. Fluorescence of single cells was assessed by Nikon Ti inverted epifluorescent microscope before performing MAPA or PF-QNM (see below) phenotyping.

### Collection of MAPA spectra

All samples were probed in phosphate buffered saline (PBS), at the room temperature. The Multimode Nanoscope IIIa atomic force microscope with a scanner E (Bruker Inc., Santa Barbara, CA) was set to oscillation (tapping) mode. SNL (Sharp Nitride Lever) probes (Bruker Inc., Santa Barbara, CA) were used, with cantilevers of 0.12 Newton/m (model substrates) or 0.06 N/m (cells and cell-bearing substrates) spring constants. The probes were tuned to near-resonant fundamental frequency in liquid (∼ 9 kHz) and engaged with the cell or a synthetic surface. The force curve of probe-surface interactions was collected and the Z-position of probe was constantly monitored to assure that engagement is not “false” and that only VDW are present with no curve hysteresis and no indentation (“forbidden zone”). The scanning was briefly carried on at 1 Hz, for areas ranging from 0.01 x 0.01µm (cells; practically point-probing) to 3 x 3 µm (model substrates). Then, the sweep of 0 to up to 800 kHz frequencies was applied to the probe and the “engaged” frequency spectra were collected at 512 points electronic resolution. After return to scanning and checking proper VDW probe engagement (Z position), the probe was lifted about 40 µm and the “non-engaged” reference spectra were collected. The target amplitude of 1V was maintained through the collections. At least three spectra were collected for each cell or model surface (technical repeats), and at least three experimental repeats were performed.

In the case of C4-2, U937 and CWC cells the same cell preparations were mechanically phenotyped with indentation-based FS-AFM using PeakForce Quantitative NanoImaging (PF-QNM) mode. Three mechanical parameters: viscoelastic adhesion, stiffness (opposite to elasticity) and deformation, were collected with Bioscope Catalyst AFM, using ScanAssist in Air probes (all from Bruker Inc., Santa Barbara, CA), as previously described^12^.

### Spectral analysis of biological surfaces using MAPA

Numerical values for amplitude as a function of frequency (frequency spectra) were used for spectral analysis. The purpose is classification of spectra was to extract their key features that allow identifying unique types of surface composition characteristic for particular classes of model substrates or cells.

The core steps to perform spectral classification^29^.

1. Spectral Preprocessing: Raw spectra were baseline corrected and denoised with Savitzky-Golay smoothing^30^.
2. Spectra collected from the same cell were averaged after testing their similarity with the Pearson correlation test or PCA.^31^
3. To limit potential overfitting in high resolution spectra, we used top principal components with the input layer after performed features reduction with Principal Component Analysis. ^32^
4. We used a Neural Network trained via Resilient Backpropagation with Backtracking to classify the processed spectra. This approach is characterized by a low risk of overshooting and the best spectral performance. ^33^
5. We performed the neural network using batch training to calculate full-batch gradient signs over the entire dataset.^34^
6. To assess the network performance we applied the confusion matrix and F1 scores.
7. Confidence limits were represented by softmax probabilities, whereas confusion matrix detected misclassifications between the highly similar spectra.^35^
8. To find which wavelengths drove the classification decisions, we applied weight analysis. Four spectral clusters were identified as shown in **Fig. 1G**.
9. To map, recognize, and reconstruct biological surfaces at nanoscale, we applied Graph Neural Networks (GNNs).
10. We directly used GNN to build a labeled surface graph, connecting nodes (structural segments) with edges (VDW bonds) in **Fig. 1H**.^36^
11. This approach overcomes physical limits of tip-sample convolution, allows for fast automated classification, and is less sensitive to noisy experimental data.
12. **Fig. 1H** represents the first step in a mesh structure analysis to reconstruct a VDW surface based on extracted spectral features of 9 distinct types of samples.
13. We constructed an SVM Territorial Map to a visually represent results of a Support Vector Machine classifier splitting a continuous 2D space into distinct territories (**Fig. 2C**). Decision Territories are colored zones representing the predicted class for any coordinate in that space. Decision Boundaries are lines separating the colored regions, where the model is exactly neutral. Scattered Data Points plotted within the territories show the original training or testing data.^37^
14. The map shows only a few misclassified points (e.g. yellow points in the green territory) and no boundary twisting that would indicate overfitting.^38^

## Acknowledgements

The work was supported by IMAT R61 CA291213 (PAO, MG).

## Notes

### Competing Interest Statement

The authors have declared no competing interest.

## References

1. McGrail DJ, McAndrews KM, Dawson MR. Biomechanical analysis predicts decreased human mesenchymal stem cell function before molecular differences. Exp Cell Res [Internet]. 2013 Mar;319(5):684–96. Available from: http://www.ncbi.nlm.nih.gov/pubmed/23228958

2. Cross SE, Jin Y-S, Rao J, Gimzewski JK. Applicability of AFM in cancer detection. Nat Nanotechnol [Internet]. 2009;4(2):72–3. Available from: http://www.scopus.com/inward/record.url?eid=2-s2.0-59849085988&partnerID=40&md5=fde4fc7ee43ee85750b7577db5cc4d85

3. Lekka M, Laidler P. Applicability of AFM in cancer detection. Nat Nanotechnol [Internet]. 2009 Feb;4(2):72; author reply 72-3. Available from: http://www.ncbi.nlm.nih.gov/pubmed/19197298

4. Lekka M, Laidler P, Gil D, Lekki J, Stachura Z, Hrynkiewicz AZ. Elasticity of normal and cancerous human bladder cells studied by scanning force microscopy. Eur Biophys J [Internet]. 1999;28(4):312–6. Available from: http://www.scopus.com/inward/record.url?eid=2-s2.0-0033049359&partnerID=40&md5=6510c8e53acdf19b7e9f90afe3e68b14

5. Lekka M, Pogoda K, Gostek J, Klymenko O, Prauzner-Bechcicki S, Wiltowska-Zuber J, et al. Cancer cell recognition - Mechanical phenotype [Internet]. 2012. Available from: http://www.scopus.com/inward/record.url?eid=2-s2.0-84858254502&partnerID=40&md5=05ab0f469a5a78279ceb335287baac79

6. D’Urso A, Purrello R, Cunsolo A, Milardi D, Fattorusso C, Persico M, et al. Electronic Circular Dichroism Detects Conformational Changes Associated with Proteasome Gating Confirmed Using AFM Imaging. Biomolecules [Internet]. 2023 Apr 20;13(4):704. Available from: https://www.mdpi.com/2218-273X/13/4/704

7. Brochard-Wyart F, de Gennes PG. Adhesion induced by mobile binders: Dynamics. Proc Natl Acad Sci [Internet]. 2002 Jun 11;99(12):7854–9. Available from: https://pnas.org/doi/full/10.1073/pnas.112221299

8. Lorenz B, Álvarez de Cienfuegos L, Oelkers M, Kriemen E, Brand C, Stephan M, et al. Model System for Cell Adhesion Mediated by Weak Carbohydrate–Carbohydrate Interactions. J Am Chem Soc [Internet]. 2012 Feb 22;134(7):3326–9. Available from: https://pubs.acs.org/doi/10.1021/ja210304j

9. Metwally S, Ferraris S, Spriano S, Krysiak ZJ, Kaniuk Ł, Marzec MM, et al. Surface potential and roughness controlled cell adhesion and collagen formation in electrospun PCL fibers for bone regeneration. Mater Des [Internet]. 2020 Sep;194:108915. Available from: https://linkinghub.elsevier.com/retrieve/pii/S0264127520304494

10. Rotsch C, Braet F, Wisse E, Radmacher M. AFM imaging and elasticity measurements on living rat liver macrophages. Cell Biol Int [Internet]. 1997;21(11):685–96. Available from: http://www.scopus.com/inward/record.url?eid=2-s2.0-0031271260&partnerID=40&md5=e2a45b0178a194590a10d5eca5c3a45c

11. Huang G, Osmulski PA, Bouamar H, Mahalingam D, Lin C-L, Liss MA, et al. TGF-β signal rewiring sustains epithelial-mesenchymal transition of circulating tumor cells in prostate cancer xenograft hosts. Oncotarget. 2016 Oct;

12. Osmulski PA, Cunsolo A, Chen M, Qian Y, Lin C-L, Hung C-N, et al. Contacts with Macrophages Promote an Aggressive Nanomechanical Phenotype of Circulating Tumor Cells in Prostate Cancer. Cancer Res [Internet]. 2021 Aug 1;81(15):4110–23. Available from: http://cancerres.aacrjournals.org/lookup/doi/10.1158/0008-5472.CAN-20-3595

13. Aceto N, Bardia A, Miyamoto DT, Donaldson MC, Wittner BS, Spencer JA, et al. Circulating Tumor Cell Clusters Are Oligoclonal Precursors of Breast Cancer Metastasis. Cell [Internet]. 2014 Aug;158(5):1110–22. Available from: http://www.scopus.com/inward/record.url?eid=2-s2.0-84907342688&partnerID=40&md5=13d397616dc2790aa8f798347869f1db

14. Ungai-Salánki R, Peter B, Gerecsei T, Orgovan N, Horvath R, Szabó B. A practical review on the measurement tools for cellular adhesion force. Adv Colloid Interface Sci [Internet]. 2019 Jul;269:309–33. Available from: https://linkinghub.elsevier.com/retrieve/pii/S0001868619300260

15. van der Waals forces. In: The IUPAC Compendium of Chemical Terminology [Internet]. Research Triangle Park, NC: International Union of Pure and Applied Chemistry (IUPAC); 2014. Available from: https://goldbook.iupac.org/terms/view/V06597

16. Chou C-W, Hung C-N, Chiu CH-L, Tan X, Chen M, Chen C-C, et al. Phagocytosis-initiated tumor hybrid cells acquire a c-Myc-mediated quasi-polarization state for immunoevasion and distant dissemination. Nat Commun [Internet]. 2023 Oct 17;14(1):6569. Available from: https://www.nature.com/articles/s41467-023-42303-5

17. Gaikwad RM, Dokukin ME, Swaminathan Iyer K, Woodworth CD, Volkov DO, Sokolov I. Detection of cancerous cervical cells using physical adhesion of fluorescent silica particles and centripetal force. Analyst [Internet]. 2011;136(7):1502–6. Available from: http://www.scopus.com/inward/record.url?eid=2-s2.0-79952597423&partnerID=40&md5=a3ba8e7da91d7b7593e607ad9843a24f

18. Osmulski P, Mahalingam D, Gaczynska ME, Liu J, Huang S, Horning AM, et al. Nanomechanical biomarkers of single circulating tumor cells for detection of castration resistant prostate cancer. Prostate [Internet]. 2014 Sep;74(13):1297–307. Available from: http://www.scopus.com/inward/record.url?eid=2-s2.0-84905898997&partnerID=40&md5=f77404ea1d265bbee1ed364eeeb50f13

19. Mahalingam D, Osmulski P, Wang C-M, Horning AM, Louie AD, Lin C-L, et al. Single-Cell Molecular Profiles and Biophysical Assessment of Circulating Tumor Cells. In: Circulating Tumor Cells [Internet]. John Wiley & Sons, Inc; 2016. p. 329–50. Available from: 10.1002/9781119244554.ch16

20. Piacenti AR, Adam C, Hawkins N, Wagner R, Seifert J, Taniguchi Y, et al. Nanoscale rheology: Dynamic Mechanical Analysis over a broad and continuous frequency range using Photothermal Actuation Atomic Force Microscopy. Arxiv. 2023;2307(16844):1–29.

21. Hsu Y-T, Osmulski P, Wang Y, Huang Y-W, Liu L, Ruan J, et al. EpCAM-Regulated Transcription Exerts Influences on Nanomechanical Properties of Endometrial Cancer Cells That Promote Epithelial-to-Mesenchymal Transition. Cancer Res [Internet]. 2016 Nov 1;76(21):6171– Available from: http://cancerres.aacrjournals.org/content/early/2016/08/24/0008-5472.CAN-16-0752.abstract

22. Garcia R, Herruzo ET. The emergence of multifrequency force microscopy. Nat Nanotechnol [Internet]. 2012;7(4):217–26. Available from: http://www.scopus.com/inward/record.url?eid=2-s2.0-84859578492&partnerID=40&md5=daa67102c8ed3f8fc3cf43e76e5917a8

23. Lü J, Yang J, Dong M, Sahin O. Nanomechanical spectroscopy of synthetic and biological membranes. Nanoscale [Internet]. 2014;6(13):7604–8. Available from: http://www.scopus.com/inward/record.url?eid=2-s2.0-84902449316&partnerID=40&md5=cd54698f8f501acf4f04d9ee715974c7

24. Canovic EP, Qing B, Mijailovic AS, Jagielska A, Whitfield MJ, Kelly E, et al. Characterizing multiscale mechanical properties of brain tissue using atomic force microscopy, impact indentation, and rheometry. J Vis Exp [Internet]. Department of Materials Science and Engineering, Massachusetts Institute of Technology, United States; 2016;2016(115). Available from: https://www.scopus.com/inward/record.uri?eid=2-s2.0-84989360009&partnerID=40&md5=b83a0f5137e57cba4df2a10f79546ef8

25. Sahin O, Quate CF, Solgaard O, Atalar A. Resonant harmonic response in tapping-mode atomic force microscopy. Phys Rev B [Internet]. 2004 Apr 28;69(16):165416. Available from: https://link.aps.org/doi/10.1103/PhysRevB.69.165416

26. Dokukin ME, Sokolov I. Nanoscale compositional mapping of cells, tissues, and polymers with ringing mode of atomic force microscopy. Sci Rep [Internet]. 2017;7(1):11828. Available from: 10.1038/s41598-017-12032-z

27. Makarova N, Peng B, Peerzade SAM, Dokukin M, Sokolov I. Imaging of Molecular Coating on Nanoparticle Surface Using AFM Ringing Mode. Microsc Microanal [Internet]. 2020 Aug 30;26(S2):3136–8. Available from: https://academic.oup.com/mam/article/26/S2/3136/6893813

28. Attwood S, Choi Y, Leonenko Z. Preparation of DOPC and DPPC Supported Planar Lipid Bilayers for Atomic Force Microscopy and Atomic Force Spectroscopy. Int J Mol Sci [Internet]. 2013 Feb 6;14(2):3514–39. Available from: https://www.mdpi.com/1422-0067/14/2/3514

29. Singh Y, Himeur Y, Farrelly CM, Kem-Meka Tiotsop Kadzue P, Rozenblit JZ, Kunovac A, et al. Graph Neural Networks for Medical Imaging Analysis and Biological Data: Integrating Topology, Geometry, Radiomics, and Generative AI. Bioengineering [Internet]. 2026 May 29;13(6):638. Available from: https://www.mdpi.com/2306-5354/13/6/638

30. V. D. dos Santos AC, Hondl N, Ramos-Garcia V, Kuligowski J, Lendl B, Ramer G. AFM-IR for Nanoscale Chemical Characterization in Life Sciences: Recent Developments and Future Directions. ACS Meas Sci Au [Internet]. American Chemical Society; 2023 Jun 16; Available from: 10.1021/acsmeasuresciau.3c00010

31. Karl Pearson. On Lines and Planes of Closest Fit to Systems of Points in Space. Philos Mag. 1901;2:559–72.

32. Beattie JR, Esmonde-White FWL. Exploration of Principal Component Analysis: Deriving Principal Component Analysis Visually Using Spectra. Appl Spectrosc [Internet]. 2021 Apr 22;75(4):361–75. Available from: https://journals.sagepub.com/doi/10.1177/0003702820987847

33. Riedmiller M, Braun H. A direct adaptive method for faster backpropagation learning: the RPROP algorithm. In: IEEE International Conference on Neural Networks [Internet]. IEEE; 1993. p. 586–91. Available from: http://ieeexplore.ieee.org/document/298623/

34. Riedmiller M. Rprop — Description and Implementation Details. Technical Report. Tech Rep. 1994;

35. Wright LG, Onodera T, Stein MM, Wang T, Schachter DT, Hu Z, et al. Deep physical neural networks trained with backpropagation. Nature [Internet]. 2022 Jan 27;601(7894):549–55. Available from: https://www.nature.com/articles/s41586-021-04223-6

36. Lanteri F, Cremonesi M. A mesh-based Graph Neural Network approach for surrogate modeling of Lagrangian free surface fluid flows. Comput Fluids [Internet]. 2025 Oct;301:106773. Available from: https://linkinghub.elsevier.com/retrieve/pii/S0045793025002336

37. Oinonen N, Kurki L, Ilin A, Foster AS. Molecule graph reconstruction from atomic force microscope images with machine learning. MRS Bull [Internet]. 2022 Sep 12;47(9):895–905. Available from: https://link.springer.com/10.1557/s43577-022-00324-3

38. Schuetzke J, Szymanski NJ, Reischl M. Validating neural networks for spectroscopic classification on a universal synthetic dataset. npj Comput Mater. 2023 Jun;9(1):100.

